# Uncoupling of sgRNAs from their associated barcodes during PCR amplification of combinatorial CRISPR screens

**DOI:** 10.1101/265686

**Authors:** Mudra Hegde, Christine Strand, Ruth E. Hanna, John G. Doench

## Abstract

Many implementations of pooled screens in mammalian cells rely on linking an element of interest to a barcode, with the latter subsequently quantitated by next generation sequencing. However, substantial uncoupling between these paired elements during lentiviral production has been reported, especially as the distance between elements increases. We detail that PCR amplification is another major source of uncoupling, and becomes more pronounced with increased amounts of DNA template molecules and PCR cycles. To lessen uncoupling in systems that use paired elements for detection, we recommend minimizing the distance between elements, using low and equal template DNA inputs for plasmid and genomic DNA during PCR, and minimizing the number of PCR cycles. We also present a vector design for conducting combinatorial CRISPR screens that enables accurate barcode-based detection with a single short sequencing read and minimal uncoupling.

## INTRODUCTION

The development and integration of oligonucleotide synthesis techniques, lentiviral vectors, and massively-parallel next-generation sequencing - the ability to write, deliver, and read DNA sequences - has enabled functional annotation of genetic elements at scale across many biological systems. Massively-parallel reporter assays (MPRA)^1,2^, genome-wide screens utilizing CRISPR technology^3^, and single-cell RNA sequencing studies^4-6^ are just some examples of experimental approaches that have employed this general framework. Often, a barcode is linked to a sequence element of interest, and thus it is imperative to understand and minimize potential sources of false calls, that is, the uncoupling of the element from its intended barcode.

It has previously been reported that barcodes used to identify open reading frames (ORFs) can uncouple from the associated ORF during the process of lentiviral production and infection, a requisite step for most pooled screening strategies^7^. Furthermore, vectors used for single-cell RNA sequencing of CRISPR screens have recently been reported to undergo similar uncoupling between the single guide RNA (sgRNA) and its associated barcode^8-10^. Other assays that rely on barcodes are also susceptible to uncoupling. In MPRA, for example, promoter or enhancer variants are typically tagged with a transcribed barcode, which is then used to infer the identity of the variant that led to expression changes^11^. Similarly, screening approaches that use unique molecular identifiers (UMIs) to obtain an absolute count of cells receiving a perturbation such as an sgRNA may be susceptible to uncoupling between the UMI and the sgRNA, potentially leading to an inflated estimate of diversity^12,13^. Recently, numerous approaches to combinatorial CRISPR screens have been described, for which accurate quantitation of two unique sgRNA sequences in the same vector presents the same challenge^14-18^.

## RESULTS

We recently developed a combinatorial screening approach, dubbed “Big Papi,” which uses orthologous Cas9 enzymes from *S. aureus* and *S. pyogenes* to achieve combinatorial genetic perturbations in pooled screens^17^. Cells that already express *S. pyogenes* Cas9 (SpCas9) are transduced with a single Big Papi vector, which delivers *S. aureus* Cas9 (SaCas9) and both an SpCas9 sgRNA and an SaCas9 sgRNA. In our original implementation, the two sgRNAs were separated by ~200 nucleotides (nts), such that both could be read out with a single sequencing read, albeit a relatively long and thus more expensive sequencing run. In order to increase the cost effectiveness of the method, we set out to reduce the required read length by incorporating barcodes into the oligonucleotides used to create these pooled libraries. However, given concerns of uncoupling, we sought to examine the fidelity of our barcoding system.

We incorporated a six nucleotide barcode into each of the sgRNA-containing oligonucleotides, immediately adjacent to the complementary regions at the 3’ end of each oligonucleotide necessary for overlap extension (Figure 1). This design places the barcodes 17 nts apart and thus requires a read length of only 29 nts to determine the combination of sgRNAs. To test the frequency of barcode uncoupling with this design, we synthesized 2 sets of 57 oligonucleotides, one for SpCas9 and one for SaCas9. To create a pooled library, we would normally mix together all the oligonucleotides to create 57 × 57 = 3,249 combinations, thus performing one pooled overlap extension reaction. Here, however, only oligonucleotides from analogous wells were mixed together - e.g. well A1 oligonucleotides for SpCas9 and SaCas9 were mixed together, etc. - for a total of 57 combinations, and 57 overlap extension reactions were performed in parallel. The resulting dsDNA products were pooled and cloned into the pPapi vector by Golden Gate cloning (see Methods). This library is thus sensitive to both uncoupling of barcodes from their associated sgRNAs, as well as to unintended combinations of sgRNAs, as only a small fraction (57 ÷ 3,249 = 1.7%) of all potential SaCas9/SpCas9 sgRNA combinations should be present.

**Figure 1.**
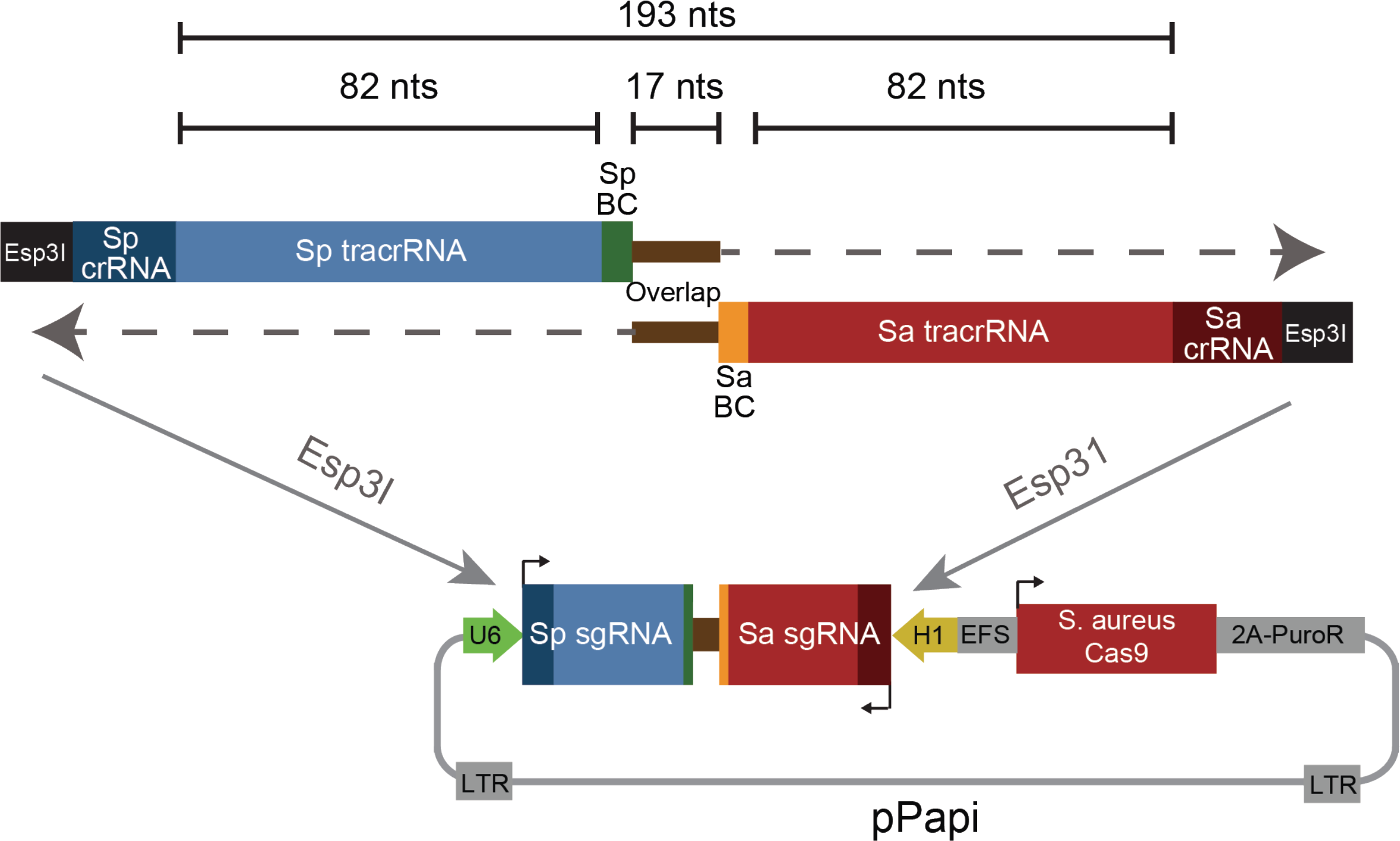
Schematic of oligonucleotide design and combinatorial library cloning strategy proposed in this study. BC: barcode.

From the plasmid DNA (pDNA) library, we generated lentivirus and infected it into A375 cells expressing SpCas9. One week after infection, sufficient time to allow any residual pDNA carried over from the production of lentivirus to degrade and dilute^7^, we prepared genomic DNA (gDNA). We then performed PCR with 28 cycles for both the pDNA (10 ng input) and gDNA (10 μg input), using primers that amplified both sgRNAs and their associated barcodes. We sequenced the resulting products with a single end read of sufficient length (300 nts) to capture all relevant sequences.

We analyzed the sequencing reads for evidence of uncoupling between sgRNAs (e.g. an SpCas9 sgRNA from well A1 appearing in combination with an SaCas9 sgRNA from any other well). We found substantially more uncoupling in the pDNA sample than in the gDNA sample, with only 64% of sgRNAs appearing with their correctly-matched sgRNA for the pDNA sample whereas 82% were correctly paired in the gDNA sample (Figure 2A). Likewise, we examined uncoupling between sgRNAs and their associated barcodes and observed that, across the 57 sgRNAs for each Cas9, a median of 79% and 92% of sgRNAs were appropriately coupled to their barcodes in the pDNA and gDNA samples, respectively (Figure 2B). These results, whereby the pDNA showed more extensive uncoupling than the gDNA, were unexpected, as only the gDNA sample had been packaged into lentivirus and integrated into cells, the steps previously suggested to generate uncoupling^7,19^. Moreover, the same pDNA had been used to generate the lentivirus infected into cells, suggesting that the pDNA uncoupling had not occurred prior to lentiviral production.

**Figure 2.**
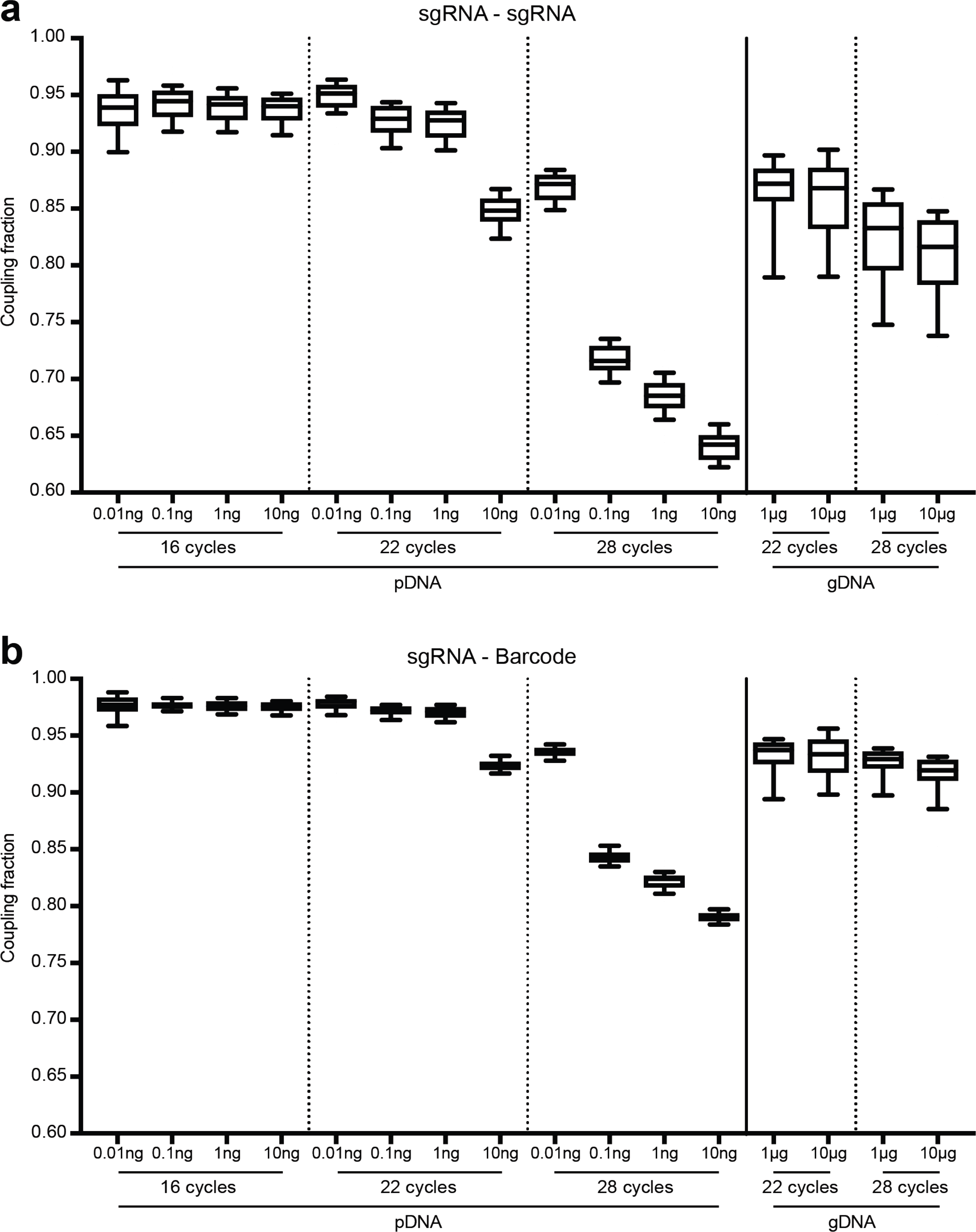
Comparison of uncoupling across sample types, input amounts, and number of PCR cycles, (a) Uncoupling between SaCas9 sgRNAs and SpCas9 sgRNAs under various PCR conditions. Each box represents 57 paired sgRNAs, plotting the fraction of reads for which the sgRNAs were correctly paired. The line represents the median, the box the 25% and 75% percentiles, and the whiskers the 10th and 90th percentiles. Our initial PCR conditions (28 cycles with 10 ng pDNA and 10 pg gDNA) led to substantial uncoupling, (b) Uncoupling between sgRNAs and their associated barcodes under various PCR conditions. Box and whisker plots are the same as In (a).

We noted that one potentially relevant difference between the two samples was the number of template molecules: 10 ng of pDNA contains ~500-fold more template molecules than 10 μg of gDNA (8.1×10^8^ vs. 1.5×10^6^ template molecules, respectively; see Methods for calculations). We also considered that the number of PCR cycles could affect uncoupling. Thus, we asked whether starting with comparable numbers of template molecules or varying the number of PCR cycles could alter the observed rates of uncoupling.

In both pDNA and gDNA samples, we found that decreasing both the number of cycles and template molecules decreased uncoupling. When using 22 cycles of PCR and approximately equal numbers of template molecules (10 pg pDNA, 10 μg gDNA), we observed that 95% and 87% of sgRNAs were correctly coupled, respectively (Figure 2A). Likewise, under these PCR conditions, a median of 98% of reads showed appropriate coupling of sgRNAs and their associated barcodes in the pDNA sample, whereas the gDNA showed 93% correct coupling (Figure 2B). Thus, when the amounts of template were normalized, the results were consistent with some uncoupling occurring during lentiviral production, as reported by Elledge and colleagues^7^.

These results implicate the PCR step as a large source of uncoupling under conditions of either higher template amounts or cycling number. One potential mechanism to explain these observations is abortive products, in which the polymerase falls off the template after it has amplified one sgRNA (or barcode) but has not finished the product. In this scenario, the 3’ end of this abortive product would still be capable of serving as a primer in the next cycle by binding to common, intervening sequence and extending, thus coupling the initial sgRNA (or barcode) to a different, unintended sgRNA (or barcode). Such abortive products may become more common as nucleotides become more limiting, as would be the case in later cycles or with more templates of input, as more products have been formed and thus fewer free nucleotides are available. This hypothesis is also consistent with the observation that there were higher levels of uncoupling between the two sgRNAs than between an sgRNA and its barcode; in this design, the two sgRNAs are separated by 193 nts, compared to just 84 nts between an sgRNA and its barcode, thus increasing the probability of a recombinant product.

Finally, to test whether the PCR polymerase had an effect on uncoupling, we compared several polymerases: the previously-used ExTaq, a hot-start version of ExTaq (ExTaq-HS), and Herculase. With 1 μg of gDNA as input and 22 cycles of PCR, we observed slightly decreased performance with Herculase, with a median sgRNA - barcode coupling fraction of 91%, compared to 93% and 94% for ExTaq and ExTaq-HS, respectively (Figure 3). With 10 μg of gDNA both ExTaq and ExTaq-HS showed a median coupling of 92%; Herculase failed to produce a product, as expected based on the manufacturer’s recommended amplification conditions. Given that combinatorial screens require a large number of cells and thus result in large amounts of gDNA, ExTaq, which tolerates higher amounts of gDNA in a reaction and shows little uncoupling under these conditions, remains our preferred polymerase.

**Figure 3.**
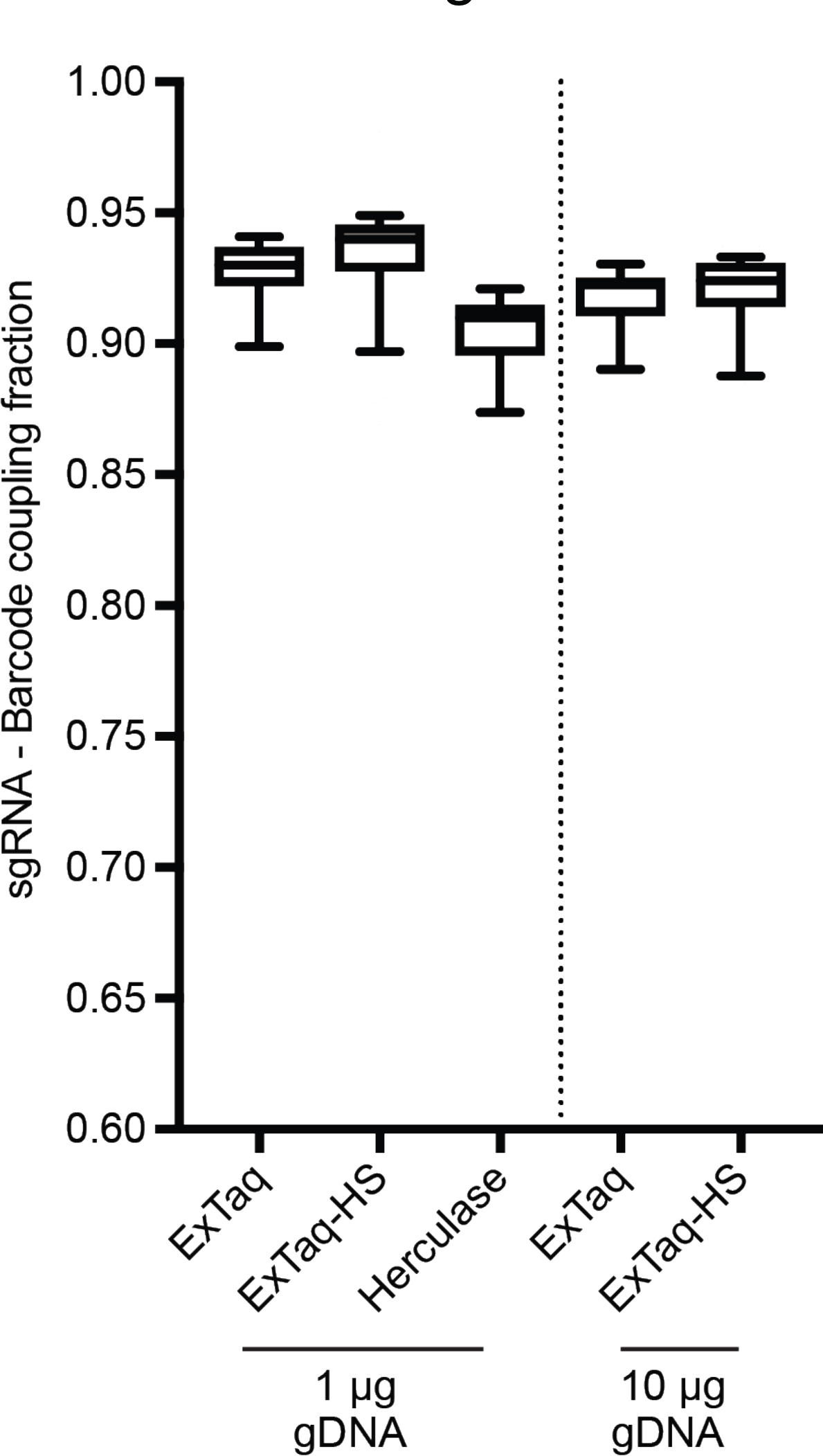
Comparison of PCR polymerases with two concentrations of gDNA. Herculase did not produce a product with 10 pg of gDNA.

## DISCUSSION

Multiple designs have been used to express pairs of sgRNAs used in combinatorial CRISPR screens (Figure 4, Table 1), and all require performing PCR to retrieve the sgRNAs or barcodes from the genomic DNA. Our results suggest that shorter distances between relevant sequences are important for alleviating PCR-based uncoupling; in current approaches, the distance between relevant elements has varied widely. Additionally, we demonstrate that minimizing the number of PCR cycles can also help mitigate uncoupling. Finally, when amplifying pDNA to serve as a measure of initial library abundance, it is important to use a similar amount of template molecules as present in the gDNA samples. The importance of PCR cycle number and template DNA input for PCR-based recombination has been previously observed^20^, but is of particular relevance given the current interest in barcode-based pooled screening.

**Table 1:**
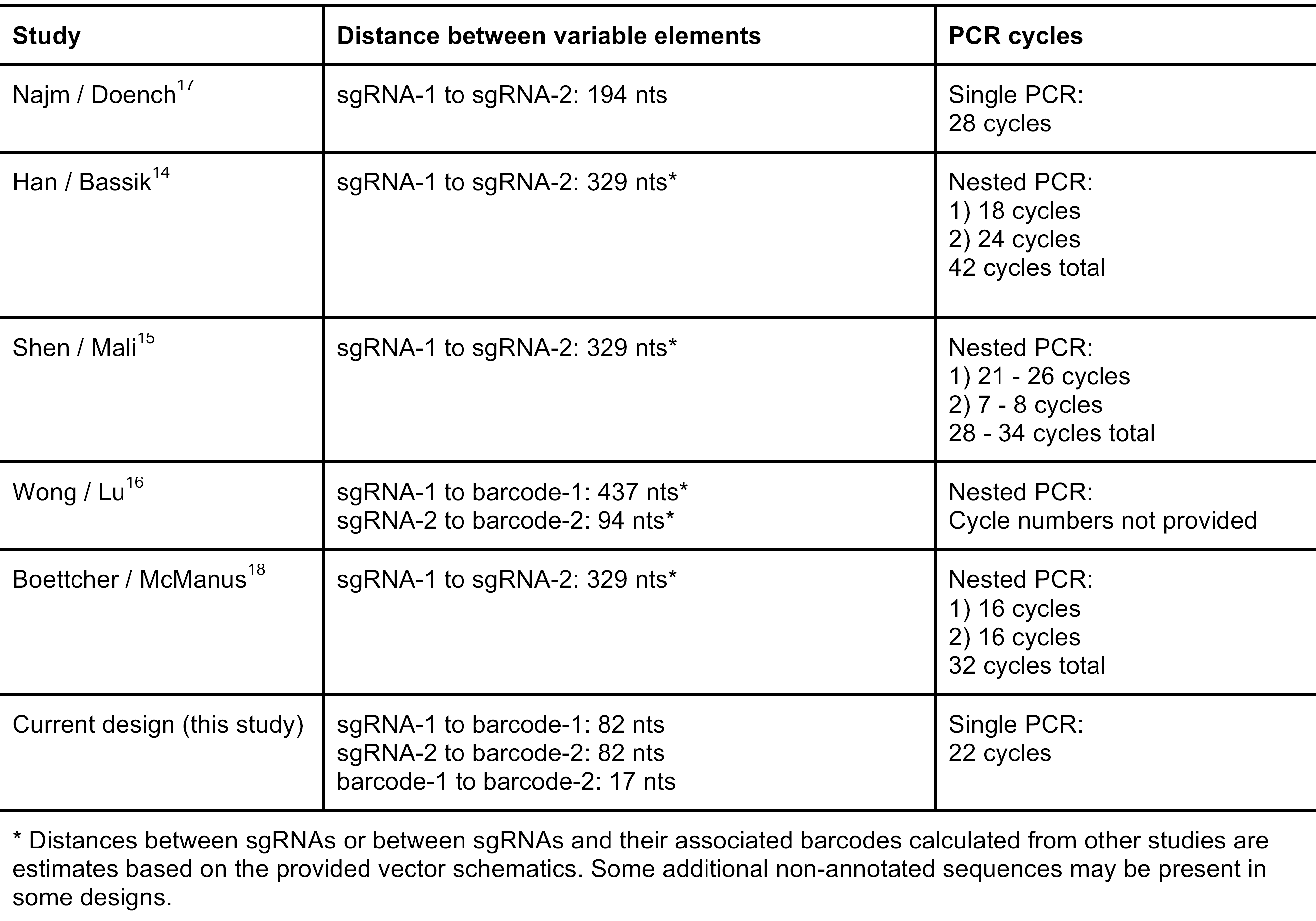

| Table 1: PCR conditions from dual-sgRNA studies |  |  |
| --- | --- | --- |
| Study | Distance between variable elements | PCR cycles |
| Najm / Doench<sup>17</sup> | sgRNA-1 to sgRNA-2: 194 nts | Single PCR:<br>28 cycles |
| Han / Bassik<sup>14</sup> | sgRNA-1 to sgRNA-2: 329 nts* | Nested PCR:<br>1) 18 cycles<br>2) 24 cycles<br>42 cycles total |
| Shen / Mali<sup>15</sup> | sgRNA-1 to sgRNA-2: 329 nts* | Nested PCR:<br>1) 21 - 26 cycles<br>2) 7 - 8 cycles<br>28 - 34 cycles total |
| Wong / Lu<sup>16</sup> | sgRNA-1 to barcode-1: 437 nts*<br>sgRNA-2 to barcode-2: 94 nts* | Nested PCR:<br>Cycle numbers not provided |
| Boettcher / McManus<sup>18</sup> | sgRNA-1 to sgRNA-2: 329 nts* | Nested PCR:<br>1) 16 cycles<br>2) 16 cycles<br>32 cycles total |
| Current design (this study) | sgRNA-1 to barcode-1: 82 nts<br>sgRNA-2 to barcode-2: 82 nts<br>barcode-1 to barcode-2: 17 nts | Single PCR:<br>22 cycles |
| * Distances between sgRNAs or between sgRNAs and their associated barcodes calculated from other studies are estimates based on the provided vector schematics. Some additional non-annotated sequences may be present in some designs. |  |  |

**Figure 4.**
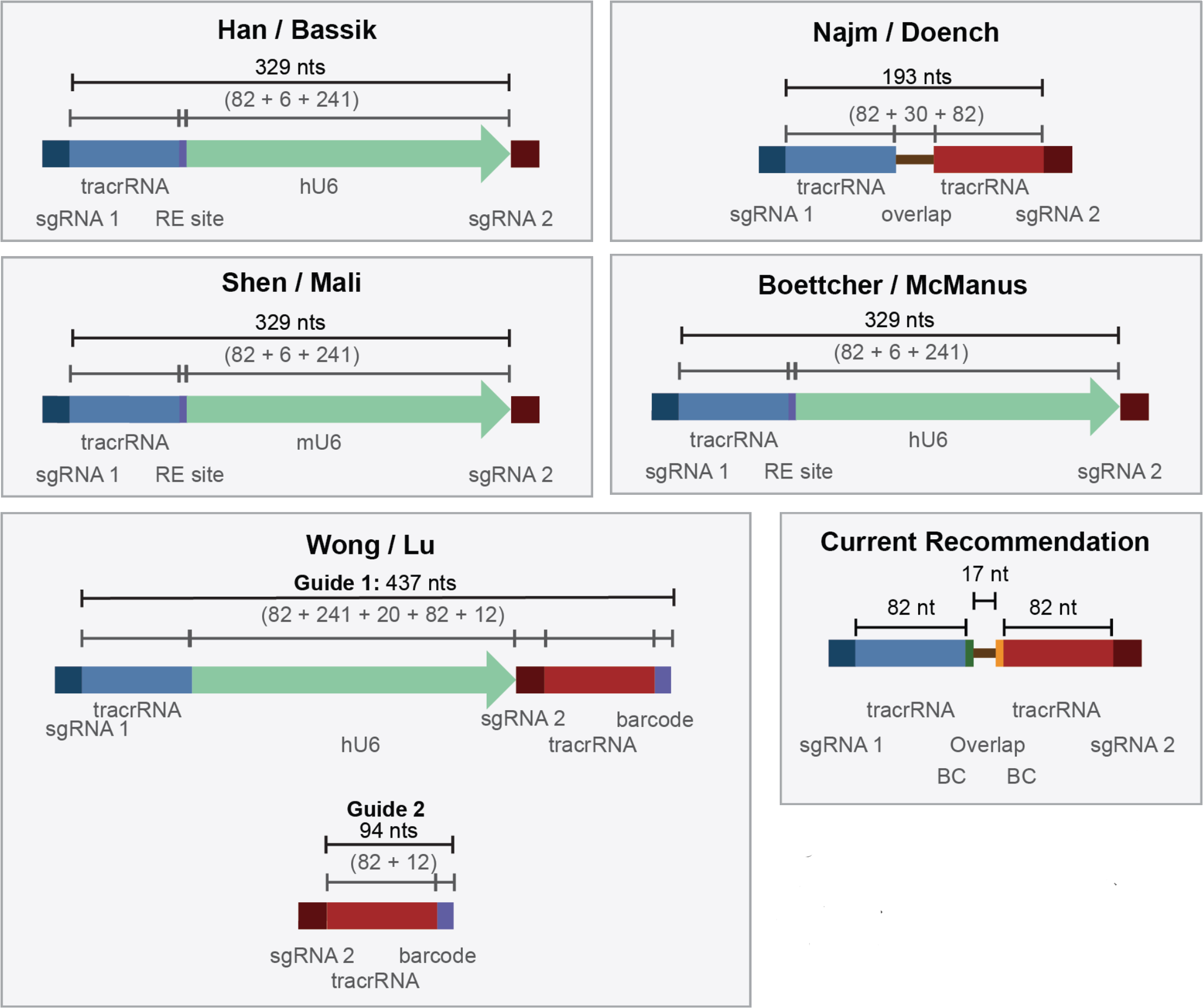
Schematics of various vector designs used for combinatorial CRISPR screens. see also Table 1.

These results also reinforce previous findings that recombination during lentiviral replication, a distance dependent factor, is another important source of uncoupling^7,19^. Thus, minimizing the distance between elements, which reduces the likelihood of uncoupling during both lentiviral replication and PCR, should be an important design parameter. Another recently proposed strategy to reduce recombination during lentiviral packaging is to dilute the library with a carrier plasmid during lentiviral production^10^, although this approach reduces viral titer by about 100-fold and thus is likely not practical for many cell-based applications.

Our current preferred combinatorial vector design has a short distance between the sgRNA and its barcode, 82 nts (the length of the tracrRNA), which results in minimal uncoupling during lentiviral production. Further, the two barcodes are only 17 nts apart, and thus there is little chance for uncoupling between barcodes during PCR retrieval of the cassette following a screen. This design should help to minimize this source of noise in combinatorial genetic screens. Additionally, these results provide guidance for optimizing many other experimental settings that use a barcode to track a sequence element of interest.

## ACKNOWLEDGEMENTS

We thank Kendall Sanson, Ellen Sukharevsky, and David Root for helpful discussions, and the entire Genetic Perturbation Platform (GPP) at the Broad Institute for being a great team. This work is supported by the Functional Genomics Consortium.

## METHODS

### Vectors

The pPapi plasmid used for dual expression of sgRNAs was previously described^17^ and is available from Addgene (#96921).

### Library Production

Two sets of oligonucleotides were ordered from Integrated DNA Technologies (IDT, Iowa). One set generates SpCas9 sgRNAs that will be expressed from the U6 promoter in pPapi, the other set generates SaCas9 sgRNAs that will be expressed from the H1 promoter. Each oligonucleotide is 139 nts in length and were ordered as Ultramers, delivered at a final concentration of 5 μ M. Oligonucleotides were mixed by well - e.g. SpCas9 A1 mixed with SaCas9 A1, SpCas9 A2 mixed with SaCas9 A2, etc. - using 2 μ L of each oligonucleotide; 6 μ L water; 10 μ L NEBnext 2× master mix (New England Biolabs M0541L). The 57 reactions were overlap-extended as follows:

a. 98°C for 3 minutes
b. 98°C for 30 seconds; 48°C for 30 seconds; 72°C for 1 minute, for 12 cycles
c. 72°C for 5 minutes

The 57 reactions were then purified by adding 5 μ L of each reaction to 1.5 mL buffer PB and proceeding with a PCR spin column purification (Qiagen 28104).

To generate pooled libraries in which combinations are not separated by individual wells, we recommend the following:

a. Pool all SpCas9 oligonucleotides at 5 μ M pool all SaCas9 oligonucleotides at 5 μ M.
b. To 10 μ L 10x ExTaq buffer and 70 uL water, add 5 μ L SpCas9 pool and 5 μ L SaCas9 pool.
c. Pre-warm heat block to 95°C, add mixture, turn off heat block, and allow to slowly cool to room temperature (~2 hours). When done, turn heat block back on as a token of good lab citizenship, although this will increase the experiment’s carbon footprint.
d. Add 8 uL dNTPs, 2 uL ExTaq (Takara RR001A), onto thermocycler: 48° for 40 minutes, 72° for 20 minutes.
e. Purify by adding to 500 μL buffer PB and proceeding with a PCR spin column purification.

The resulting dsDNA is then ligated into the BsmBI-digested pPapi vector using Golden Gate assembly:

5 μL Tango Buffer (ThermoFisher)

5 μL DTT (stored at −80°C and used once, 10 mM stock)

5 μL ATP (stored at −80°C and used once, 10 mM stock)

500 ng pPapi vector, pre-digested with Esp3I or BsmBI, gel-extracted, and isopropanol-precipitation purified

100 ng dual sgRNA dsDNA insert

1 μL Esp3I (ThermoFisher ER0452)

1 μL T7 ligase (Enzymatics, 3,000 Units / μL L6020L)

Up to 50 μL water

Cycle 100x (overnight): 5 minutes at 37°C, 5 minutes at 20°C.

Purify Golden Gate product by isopropanol precipitation. Per 50 μL reaction, add in order:

1 μL GlycoBlue (Ambion AM9515)

4 μL NaCl, 5M 55 μL isopropanol

a. Vortex, and incubate at room temperature for 15 minutes.
b. Centrifuge at >10,000g for 15 minutes at room temperature.
c. Remove liquid, avoiding the pellet (it is okay to leave a little liquid behind).
d. Add 950 μL 70% EtOH, vortex, centrifuge for 5 minutes at room temperature, remove liquid.
e. Repeat step (c).
f. Centrifuge for 1 minute and remove any residual liquid with a fine-tipped pipette (e.g. P200 or smaller); allow to air dry for 1 minute.
g. Resuspend with 10 μL water or TE, on ice. Flick the tube and briefly centrifuge as needed.

To transform the library into *E. coli,* we recommend STBL4 cells (Invitrogen 11635018). Add 10 μL of isopropanol-precipitated DNA to 100 μL electrocompetent cells. This step will need to be scaled as library size increases.

### Virus Production

Pooled library virus was made using the same large scale T175 flask method used previously^17^. Briefly, 24 hours pre-transfection, 18 × 10^6^ HEK293T cells were seeded into a 175 cm^2^ tissue culture flask with 24 mL of DMEM + 10% FBS. Next day, one solution of Opti-MEM (Corning, 6 mL) and LT1 (Mirus, 305 μL) was combined with a DNA mixture of the packaging plasmid pCMV-VSVG (Addgene 8454, 5 μg), psPA×2 (Addgene 12260, 50 μg), and sgRNA-containing vector (pPapi, 40 μg). This mixture was incubated for 20-30 min at room temperature, during which media was changed on the HEK293Ts. Following incubation, the transfection mixture was added dropwise to cells. The cells were incubated for 6-8 h, after which time media was replaced with DMEM + 10% FBS, supplemented with 1% BSA. 36 hours post-media replacement, virus was harvested.

### Cell culture

A375 cells were obtained from the Cancer Cell Line Encyclopedia. Cells were cultured in RPMI + 10% FBS, routinely tested for mycoplasma contamination and maintained in a 37 °C humidity-controlled incubator with 5.0% CO_2_. Cells were maintained in exponential phase growth by passaging every 2 or 3 days. Cell lines were maintained without antibiotics, and supplemented with 1% penicillin/streptomycin post-lentiviral infection. The A375 Cas9 derivative was made by transducing with the lentiviral vector pLX_311-Cas9, which expresses blasticidin resistance from the SV40 promoter and Cas9 from the EF1α promoter (Addgene 96924).

### Infection Optimization

A375 cells stably expressing SpCas9 were infected as described previously^17^.

### Genomic DNA Preparation

Genomic DNA (gDNA) was isolated using the QIAamp DNA Blood Maxi Kit (Qiagen) as per the manufacturer's instructions. Resulting gDNA was quantitated by UV Spectroscopy (Nanodrop).

Going forward, we recommend the use of Nucleospin Blood XL kits (Macherey-Nagel, 740950) for gDNA isolation, and the use of Qubit with the dsDNA BR kit (Invitrogen Q32850) to quantitate concentration.

### Calculations for Templates of Input

GDNA: 1 template = 1 cell (assuming 3×10^9^ basepairs per cell) = 6.6 pg gDNA 10 μg gDNA × 1 cell / 6.6 pg gDNA = 1.5×10^6^ template molecules pDNA: 1 template = 1 plasmid of 12.3 kB (7.5 × 10^6^ g/mol) = 1.24 ×10^−5^ pg pDNA 10 pg gDNA × 1 plasmid / 1.24 ×10"^5^ pg pDNA = 8.1×10^5^ template molecules

### PCR and Sequencing Methods

Dual sgRNA cassettes were PCR-amplified and barcoded with sequencing adaptors using ExTaq DNA Polymerase (Clontech). Each 100 μ L total volume reaction contained 10 μ L of 10x buffer; 8 μ L dNTP (provided with the enzyme); 0.5 μ L of P5 stagger primer mix (stock at 100 μ M concentration); 10 μ L of a uniquely barcoded P7 primer (stock at 5 μ M concentration); and 1.5 μ L polymerase enzyme (ExTaq, Takara RR001A; ExTaq-HS, Takara RR042A; Herculase, Agilent 600310-51). After adding gDNA or pDNA, any remaining volume was water.

P5/P7 primers were synthesized at Integrated DNA Technologies (IDT):

#### Forward (P5)

5’AAT GAT ACGGCGACCACCGAGAT CT ACACT CTTT CCCT ACACGACGCT CTT CCGA TCT[s]TTGTGGAAAGGACGAAAC*A*C*C*G

#### Reverse (P7)

5’CAAGCAGAAGACGGCATACGAGATNNNNNNNNGTGACTGGAGTTCAGACGTGT GCTCTTCCGATCTCCAATTCCCACTCCTTTCAA*G*A*C*C

P5/P7 flow-cell attachment sequence

Illumina sequencing primer

[Stagger region] / Barcode region

Vector primer binding sequence

* between bases indicate phosphorothioate linkages

#### PCR cycling conditions

a. 1 minute at 95°C
b. 30 seconds at 94°C, 30 seconds at 52.5°C, 30 seconds at 72 °C, for *n* cycles
c. 10 minutes extension at 72 °C.

Following PCR, samples were purified with Agencourt AMPure XP SPRI beads (Beckman Coulter A63880) according to manufacturer's instructions. Each purified pool was quantitated on with UV spectroscopy (Nanodrop) and pooled into a master sequencing pool such that each PCR well contributed approximately equally to the final master pool. The master pool was sequenced on a MiSeq sequencer (Illumina) with 300 nt single-end reads, loaded at 60% with a 5% spike-in of PhiX DNA.

### Analysis

Reads of the first sgRNA were counted by searching for CACCG, part of the vector sequence that immediately precedes the 20-nucleotide U6 promoter-driven SpCas9 sgRNA. The sgRNA sequence following this search string was mapped to a reference file with all SpCas9 sgRNAs in the library. Reads of the SpCas9 sgRNA-associated six-nucleotide barcodes were then counted by searching for part of the SpCas9 tracr sequence that precedes the barcode. The barcode was then mapped to a reference file with all SpCas9 sgRNA-associated barcodes.

Reads for the H1 promoter-driven SaCas9 sgRNA were counted by searching for part of the reverse complement of the SaCas9 tracr sequence (CTTAAAC). The 21-nucleotide sgRNA sequence following the search string was mapped to the reference file with all SaCas9 sgRNAs in the library. Reads for the six-nucleotide barcode associated with the SaCas9 sgRNA were then counted by searching for part of the overlap extension region preceding the barcode. The barcode was then mapped to the reference file with all SaCas9-associated barcodes.

